# Adoptive transfer of CTLA4-Ig-modulated dendritic cells improves TNBS-induced colitis

**DOI:** 10.1101/669630

**Authors:** Lisiery Negrini Paiatto, Fernanda Guimarães Drummond Silva, Áureo Tatsumi Yamada, Wirla Maria Silva Cunha Tamashiro, Patricia Ucelli Simioni

**Affiliations:** Department of Biomedical Science, Faculty of Americana, FAM, 13477-360, Americana, SP, Brazil; Department of Food, School of Nutrition, Federal University of Ouro Preto, 35400-000, Ouro Preto, MG, Brazil.; Department of Biochemistry and Tissue Biology, Institute of Biology, University of Campinas (UNICAMP), 13083-862, Campinas, SP, Brazil.; Department of Genetics. Evolution, Microbiology and Immunology, Institute of Biology, UNICAMP, 13083-862, Campinas, SP, Brazil.; Postgraduate Program in Biological Science (Cellular and Molecular Biology), Department of Biochemistry and Microbiology, Institute of Biosciences, Univ Estadual Paulista, UNESP, 13506-900 Rio Claro, SP, Brazil; Department of Biomedical Science, Faculty of Americana, FAM, 13477-360, Americana, SP, Brazil.

**Keywords:** colitis, dendritic cell, immune modulation, CTLA4-Ig, tolerance, inflammation, inflammatory bowel disease

## Abstract

Dendritic cells (DCs) play a crucial role in balancing immune responses, and in that sense the interactions between the B7-1 and B7-2 molecules expressed on DCs and CD28 and CTLA-4 on helper T cells are fundamental. While coupling of B7 and CD28 molecules activates immune responses, binding of B7 to CTLA4 results in its blockade. CTLA4-Ig fusion protein, a competitor molecule of the B7-CD28 interaction, has been used for the development of immunological tolerance both experimentally and in patients. Here, we evaluated the effects of adoptive transfer of bone marrow-derived dendritic cells (BMDCs) pulsed with CTLA4-Ig in TNBS-induced colitis. CTLA4-Ig-modulated BMDCs or naïve BMDC were administered intravenously to BALB/c mice prior to TNBS rectal instillation. Five days later, spleens and colon segments were removed for immunological and histological analysis. Our results showed that the adoptive transfer of CTLA4-Ig-modulated BMDCs was able to reduce the severity of inflammation caused by the administration of TNBS, in view of tissue integrity and reduced leukocyte infiltration in the colon segments of the treated mice compared to controls. Non-specific spleen cell activation in vitro showed a reduction in the frequency of CD4^+^ IL-17^+^ T cells and CD4^+^ IFN-γ^+^ T cells as well as IL-9 secretion in cultures. To our knowledge, this is the first description of the beneficial effects of treatment with CTLA4-Ig modulated BMDC in experimental colitis.

## 1. Introduction

Two distinct signals are required for the activation of the adaptive immune response. The first signal is established by the binding of antigen-specific receptors on the surface of T lymphocytes (TCR) and antigenic fragments associated with MHC molecules on the surface of antigen-presenting cells (APCs). The second signal comes from the engagement between co-stimulatory molecules, among which the CD28 expressed on the surface of T lymphocytes and B7-1 (CD80) and B7-2 (CD86) on the surface of APCs are prominent. The low affinity interaction between CD80/CD86 and CD28 is essential to promote the activation, proliferation and survival of lymphocytes. The production of specific antibodies by B lymphocytes and the increase of phagocytic cell activities are the most evident results of this initial activation (1–4).

As the immune response proceeds and the antigen triggering this response is eliminated, a cell surface glycoprotein called CTLA4 (CD152) begins to be expressed at low levels in activated T cells. CTLA-4 binds with high affinity to the costimulatory molecules CD80 and CD86 in dendritic cells (DCs), thus initiating the reduction of the specific immune response(5–9).

Altered responses against self-antigens are at the origin of autoimmune diseases. Inflammatory bowel disease (IBD) is a group of immune-mediated diseases characterized by severe inflammation of the digestive tract (10). The etiology of IBD is still unknown, but the most plausible hypothesis is that it is due to the combination of genetic and environmental factors, particularly disturbances of the microbiota, leading to an aberrant inflammatory response of the host (11–13). Several immunomodulatory drugs such as azathioprine and mycophenolate (inhibitors of T-cell proliferation), monoclonal antibodies, such as OKT3 (depletes and blocks T cells), and cyclosporine, tacrolimus and glucocorticoids (blockage of cytokine production) have been applied with relative success in the control of autoimmune diseases. However, most of them can lead to complications related to the onset of opportunistic infections as well as nephrotoxicity. Due to the serious side effects of nonspecific anti-inflammatory drugs and broad-spectrum immunosuppressive drugs routinely employed in the treatment of autoimmune diseases, current studies are looking at ways to manipulate immune system to reduce the need for these substances (10). Thus, new therapeutic approaches aimed at inhibiting immune responses in a more natural way have been developed in the last two decades. Among these new approaches, one of the best studied involves the use of CTLA4-Ig, a competitor molecule of the B7/CD28 interaction. In principle, its use would allow the development of immune tolerance to autoantigens by naturally blocking the activation of specific T lymphocytes (14–19).

To test these approaches prior to being screened in humans, several experimental models are available. In relation to IBD, experimental models of colitis induced by chemical or biological agents that mimic the main characteristics of human disease are currently used (20–22). Colitis induced by instillation of 2,4,6-trinitrobenzenesulfonic acid (TNBS) in BALB/c mice, for example, generates a relatively mild inflammation of the intestinal mucosa and slight weight loss, with reestablishment of the animal within a few days after instillation. Such characteristics make this one of the experimental models most used in the study of colitis modulators(23,24).

It is well known that DCs can interfere in the balance between immunity and tolerance. However, few clinical applications have been successful so far (25–27). On the other hand, some experimental studies have shown that CTLA4-Ig, a soluble chimeric fusion protein (CD152/Fc), can block the B7/CD28 signaling pathway by competition with CD80 and CD86 molecules expressed in DCs, thereby reducing responses autoimmune and graft rejection (16,28).

In this work, we evaluated the effects of the adoptive transfer of CTLA4-Ig-modulated bone marrow-derived dendritic cells (BMDC_CTLA4-Ig_) on the inflammatory response observed in TNBS-induced colitis in BALB/c mice. BMDC_CTLA4-Ig_ and control BMDCs (BMDC_naïve_) were administered intravenously for three consecutive days prior to instillation of TNBS. Five days later, the spleen and colon segments were removed for immunological and histological analysis.

## 2. Material and methods

### 2.1. Animals

BALB/c female mice (20–25g) at four weeks of age were obtained from the Multidisciplinary Center for Biological Research (CEMIB) of the University of Campinas (UNICAMP), Campinas, SP, Brazil. They were maintained in specific pathogen-free environment at 25° C ±1 and photoperiod of 12/12 hours. Mice were fed with autoclaved Nuvilab CR-diet (Colombo, PR, Brazil) and water *ad libitum* for 2-4 weeks before being used in experiments. Mouse manipulation were carried out in accordance with the ‘Guide for the Care and Use of Laboratory Animals’, as promoted by the Brazilian College of Animal Experimentation (COBEA) and approved by the Ethics Committee for Animal Experimentation of University of Campinas (CEUA/UNICAMP Protocol #3077-1). All experimental procedures were performed under anesthesia (ketamine and xylazine) and all efforts were made to minimize animal suffering. Each experimental group consisted of at least five animals. Assays were repeated at least two times. Mice were monitored daily for signs of colitis such as rectal swelling, rectal bleeding, soft stools as well as weight loss.

### 2.2. Bone marrow dendritic cells

Bone marrow dendritic cells (BMDCs) were generated from bone marrow precursors as described elsewhere. Briefly, bone marrow cells were flushed from femurs and tibias of naïve BALB/c mice with RPMI 1640 medium (Sigma) containing 10% fetal bovine serum (FBS, Cultilab, Brazil), 20 μg/mL gentamicin solution (Sigma). Bone marrow cells were seeded in six-well plates (Corning, USA) at a density of 2×10^6^ white cells/well in RPMI-10% FBS containing 20ng/mL mouse recombinant granulocyte macrophage colony-stimulating factor (mrGM-CSF) (Biosource, USA) and then incubated at 37° C in 5% CO_2_. On days 3 and 6, the culture medium was replaced with fresh medium containing GM-CSF(29). On the eighth day of culture, the differentiated cells were collected, pelleted by centrifugation at 200 g, for 10 min, resuspended in RPMI-10% FBS containing 40ng/mL CTLA4-Ig (CD152/Fc chimera, non-cytolytic, from mouse, Sigma C4358), plated in 24-well culture plates at a density of 2 x 10^6^ cells/well and cultured for an additional 24 hours (BMDC_CTLA4-Ig_). Cells cultured in the absence of CTLA4-Ig were used as control (BMDC_naïve_).

### 2.3. Phenotypic profile of BMDC

The phenotypic characteristics of the BMDCs were evaluated by flow cytometry as previously described by our group (30) and others (31–33)(33). For this, the cells were labeled with anti-mouse CD11c-APC (Clone: HL3), anti-mouse MHC-II-PE (130-091-368, Miltenyi Biotec), anti-mouseCD80-FITC (Clone: 16-10A1), anti-mouseCD86-FITC (Clone: GL1) and anti-mouse CD40-FITC (Clone: 3/23) according to the manufacturer’s instructions (BD Bioscience, USA). All controls were performed using irrelevant isotype staining. The readings were performed using the FACSCalibur (Becton-Dickinson, Franklin Lakes, NJ, USA) flow cytometer, using FCS Express 5 Plus, Research Edition software.

### 2.4. Adoptive transfer of BMDC and colitis induction

Three doses of 1×10^6^ BMDC_CTLA4-Ig_ or BMDC_naïve_ were injected intravenously into naïve syngeneic mice on days 5, 3 and 1 before the colitis induction (Fig 1). Colitis was induced by intrarectal administration of a single dose of 2,4,6-trinitrobenzenesulphonic acid (TNBS), as described elsewhere, with modifications. Briefly, mice were anesthetized with and instilled with 100µL of 1.0mg/mL TNBS (2,4,6-trinitrobenzenesulfonic acid; Sigma, USA) dissolved in 50% ethanol into the lumen of the colon. To ensure that TNBS enter entire colon, mice were held in a vertical position for 30s. Two control groups of mice that did not receive DC were used: 1) animals inoculated intrarectally with 100μL 50% ethanol in saline; 2) animals inoculated intrarectally with 100μL of 1.0mg/mL TNBS dissolved in 50% ethanol.

**Figure 1.**
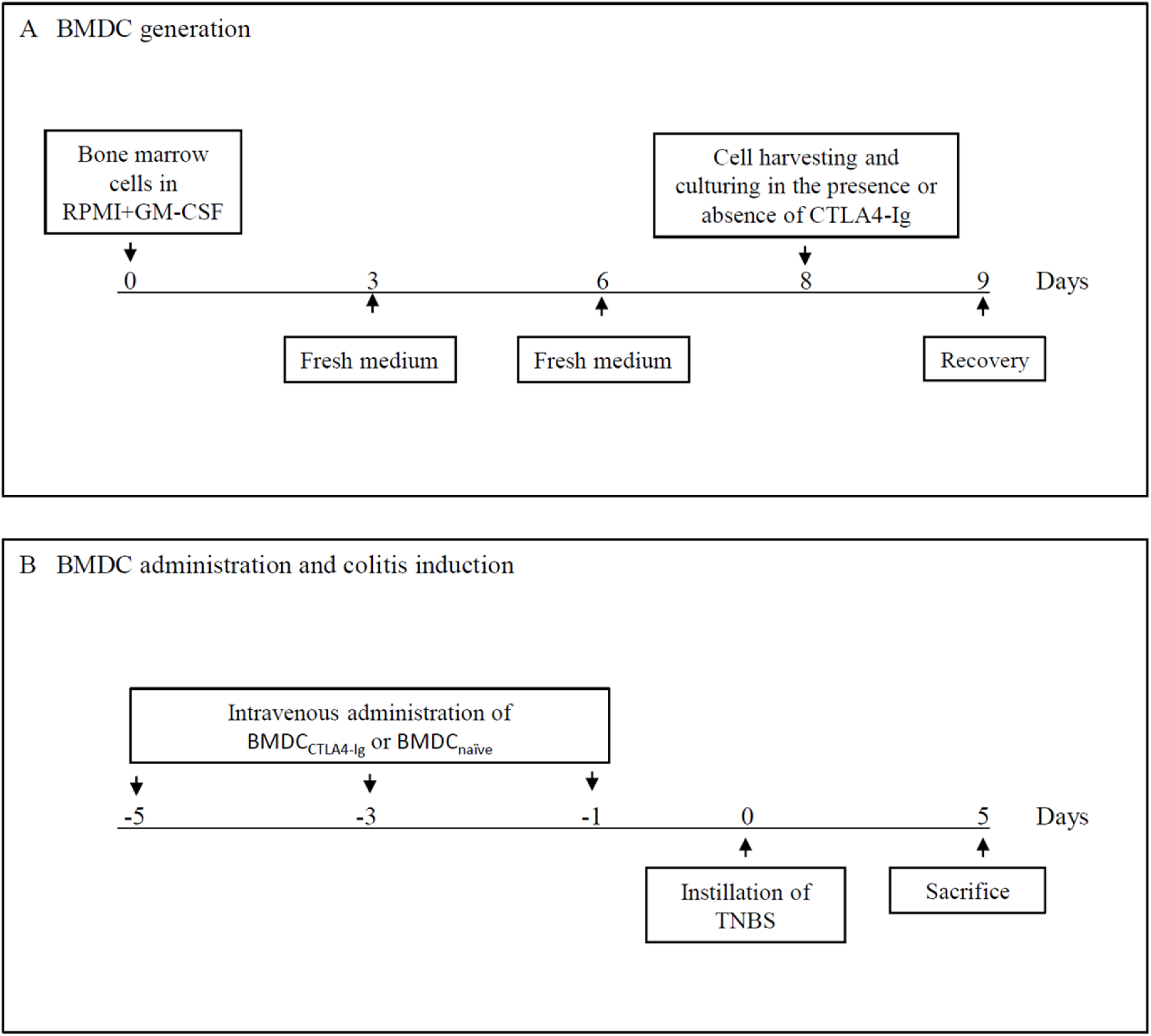

### 2.5. Evaluation of clinical symptoms of TNBS-induced colitis

Animals of all groups were weighed daily until sacrifice on the fifth day following the instillation of TNBS. Weight variation was calculated as percentage, considering the weight at day zero as 100%. Clinical symptoms such as diarrhea, rectal prolapse, bleeding and cachexia were registered and assigned as scores, ranging from 0 to 2, with 0: no change, 1: slight change (liquid feces, inflammation in the anus, mucus liberation and weakness), and 2: severe change (diarrhea, rectal prolapse, bleeding, and cachexia) and these values were mean to each animal. Data are presented as the mean ± Standard Error of the Mean (S.E.M.)

### 2.6. Histological analysis of the colon

The mice were euthanized five days after induction of colitis. Two portions of the colon distant 1-2 cm (P1 = proximal1) and 2-3 cm (P2 = proximal 2), respectively, of the anal sphincter were removed, fixed in 4% buffered formalin, dehydrated with ethanol solutions, and embedded in paraffin (Paraplast Plus Sigma P3683). Slices of 5μm were cut into a microtome (Leica - model Jung Biocut 2035) and mounted on clean glass slides. The specimens were then dewaxed, rehydrated and stained with hematoxylin and eosin (Merck). As the distal portions of the intestine are unaffected by treatment with TNBS, sections of these segments were not examined.

The P1 and P2 segments of the colon were evaluated by microscopy for the presence of folds, hemorrhage, gauge and leukocyte infiltrate. The data were represented by scores corresponding to the sum of the values attributed to the presence and characteristics of the folds in the mucosa (0 for normal folds, 1 for slightly altered folds, 2 for deformed folds, 3 for very deformed folds, 4 for dorsal folds reduction, 5 for lack of folds); bleeding (0 for no bleeding, 1 for bleeding present, 2 for large bleeding); mucosal dilatation (0 for absence of voids, 1 for apparent voids, 2 for large voids); and lymphocytic infiltrate in the mucosa, submucosa and mesentery (0-none; diffuse inflammatory infiltrate-1; 2-considerable inflammatory infiltrate with submucosal disorganization; 3-intense infiltrate) as previously described 5. Thickening of the colon wall was measured in micrometers using the Infinity Analyze Nikon H600L (100X). The final scores represent the mean ± S.E.M (23,34).

Histomorphometric analysis was performed on sections of P1 and P2 segments prepared for immunoperoxidase reactions using the following antibodies: anti-CD3 (T lymphocytes) and anti-F4/80 (macrophages) and anti-Ly-6c NIM (neutrophils) (35,36). Peroxidase-conjugated secondary antibodies (Sigma) and diaminobenzidine (DAB; Sigma) were used in the development of reactions (37,38). After counter-staining with hematoxylin/eosin, the tissues were observed under light microscopy for counting the labeled cells. Evaluations were done in a double-blind fashion and the quantification of labeled cells was performed in five random fields in each specimen. Sections P1 and P2 of at least three animals in each group were evaluated, making a total of 15 measurements per group, in each experiment. Results were expressed as mean ± S.E.M. Three independent experiments were performed (24,39,40).

### 2.7. Spleen cell proliferation

Spleens were collected aseptically from mice of all experimental groups, macerated individually, suspended in lyses buffer and pelleted by centrifugation at 200 g for 10 min. Cell concentrations were adjusted to 1×10^6^ cells/mL in RPMI medium (Sigma, USA) supplemented with 10% fetal bovine serum (Cultilab, Campinas, Brazil). After washing, spleen cell suspensions were incubated with 25μM carboxyfluoresceinsuccinimidyl probe ester (CFSE) in RPMI-10% FBS at room temperature for 5 min, according manufacturer’s recommendations (Invitrogen, USA). Cells were then pelleted by centrifugation and suspended in fresh medium. To determine the maximum uptake of CFSE, aliquots of each cell suspension were fixed with 1% formaldehyde in PBS and analyzed by flow cytometer. Then 100μL aliquots of each suspension of CFSE-labeled cells were seeded in duplicate in 96-well plates (Corning) and incubated in the presence of 2.5 μg / ml ConA for 72 hours at 37° C. Cultures of cells conducted in the absence of Con-A were used as controls.

The proliferation of T lymphocytes in the cultures was assessed at the gate of CD4+CFSE+ cells. Acquisitions were performed with FACSCalibur flow cytometer (FACSCalibur flow cytometer, BD Becton Dickinson, San Jose, CA)(34,41). The results were analyzed with the FCS Express Plus Research Edition software (FCS Express Launcher). Results were expressed as proliferation index (fold change) calculated in relation to that of the control group (24,34).

In parallel, cultures of CFSE-unlabeled spleen cells from mice of all experimental groups were conducted to measure the levels of cytokines released in the supernatants after Con-A stimulation as described below.

### 2.8. Phenotypic profile of T-cells

The frequencies of TCD4^+^CD25^+^ Foxp3^+^ (Treg cells), TCD4^+^IL17^+^, TCD4^+^IFNγ^+^ and TCD4^+^IL-10^+^ cells in the spleen cell cultures were assessed by flow cytometer. Briefly, cell suspensions were washed and initially stained with anti-CD3 APC (clone 145-2C11, BD #553066), anti-CD4-PE (Clone GK1.5) and anti-CD25-FITC (Clone 7D4). Then, cells were permeabilized by the addition of fixation/permeabilization buffer (Cytofix/Cytoperm fixation/permeabilization kit, Becton-Dickinson, BD). Suspension was stained with anti-Foxp3-APC (clone FJK-16S), anti-IL-17-APC (clone eBIO17B7) or Alexa Fluor 647 (Clone TC11-18410), anti-IFN-γ-APC (Clone XMG1.2) and IL-10-APC (Clone JESS-16E3), 647 (Clone Q21-378), according to manufacturer’s instructions. Controls were performed with irrelevant isotype staining. Acquisitions were performed with FACScalibur flow cytometer and analyzes were done with the FCS Express 5 Plus, Research Edition software (23,34)

### 2.9. Determination of Th1, Th2, Th17 and Th9 cytokines

IL-2, IL-4, IL-6, IL-10, IL-17A, IFN-γ and TNF-α were quantified in culture supernatants of spleen cells by flow cytometer, using Multiplex CBA kit (BD Cytometric Bead Array Th1/Th2/Th17, San Diego, USA) according to manufacturer’s instructions. Fluorescence were acquired in FACSCalibur cytometer and analyzed with FCAP Array TM Software Version 3.0 (BD). IL-9 determination was assayed with CBA flex set (BD Cytometric Flex Set Th9, San Diego, USA) (23,34)

### 2.11. Statistical analysis

The statistical analysis was performed using GraphPad Prism 5 (GraphPad Software, San Diego, CA, USA). The statistical significance of differences between control and experimental groups were determined by one-way ANOVA, followed by Bonferroni’s test for multiple comparisons or unpaired Student’s t-test. The results were expressed as mean ± Standard Error of the Mean (S.E.M). Values were considered significant at P< 0.05. All data presented are representative of at least three independent experiments.

## 3. Results

### The effects of adoptive transfer of BMDCs on TNBS-induced colitis

Dendritic cells were differentiated in vitro from precursors collected from the bone marrow of BALB/c mice by their culture in the presence of recombinant GM-CSF for eight days. Differentiated bone marrow DCs (BMDCs) were then incubated in the presence or absence of recombinant CTLA4-Ig for 24 hours before being employed in adoptive transfer assays. We observed that modulation of BMDCs with CTLA4-Ig did not modify the expression pattern of the CD11c, MHC class II, CD40, CD80 and CD86 molecules on the surface of these cells compared to CTLA4-Ig untreated BMDCs (BMDCnaïve), as can be seen in Supplementary Figure 1 (Supp. Fig 1).

Weight loss in TNBS-induced colitis is generally mild (about 10%) and recovery of body weight is usually observed as early as the fourth day after drug administration in BALB/c mice. As can be seen in Figure 2A, the adoptive transfers of BMDC_CTLA4-Ig_ or BMDC_naïve_ did not result in any significant improvement in weight loss observed in the first days after TNBS instillation and, as expected, by the fourth day all mice had already recovered. However, the other clinical signs of the disease (diarrhea, rectal prolapse, soft stools, and hemorrhagic stools) were significantly reduced by previous treatment with BMDC_CTLA4-Ig_, as shown in Figure 2B.

**Figure 2:**
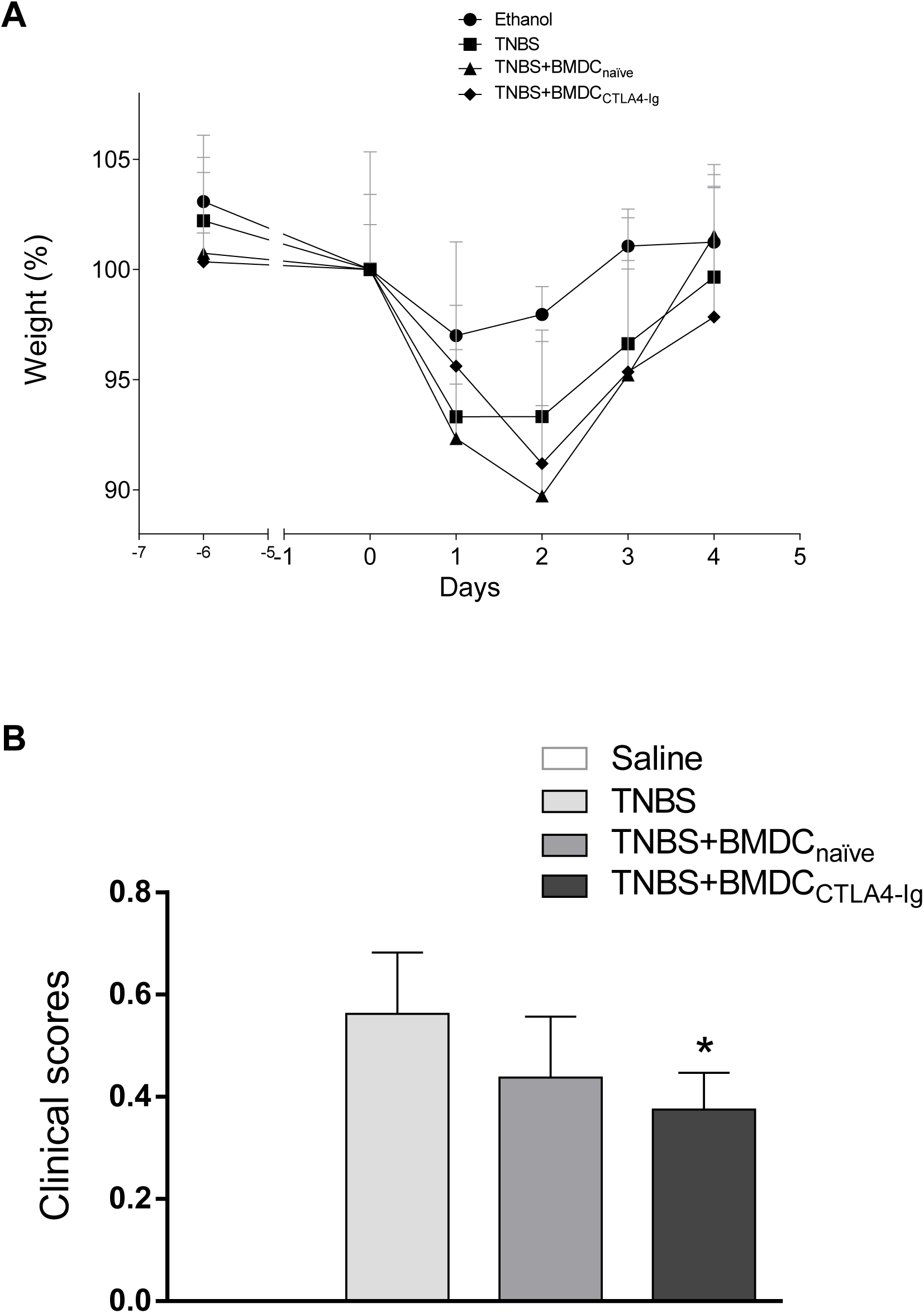

As can be seen in Figure 3A, administration of TNBS induced a strong inflammatory reaction that affected the regions closest to the rectum (both P1 and P2 segments), with a large number of infiltrated leukocytes. However, previous treatment with BMDC_CTLA4-Ig_ significantly reduced leukocyte infiltration caused by instillation of TNBS.

The cells present in the infiltrate consisted mainly of macrophages (Figure 3B), while neutrophils were rarely found. It was also observed that the colonic tissue of mice treated with BMDC_CTLA4-Ig_ showed a significant reduction in the number of infiltrated macrophages when compared to the tissues of mice without previous treatment with BMDCs. Pretreatment with BMDC_naïve_ did not modify the inflammatory process induced by the instillation of TNBS. On the other hand, the adoptive transfer of BMDC_CTLA4-Ig_, but not BMDC_naïve_, was able to prevent thickening of the colon wall, particularly in the P1 region, as shown in Figure 3C.

**Figure 3:**
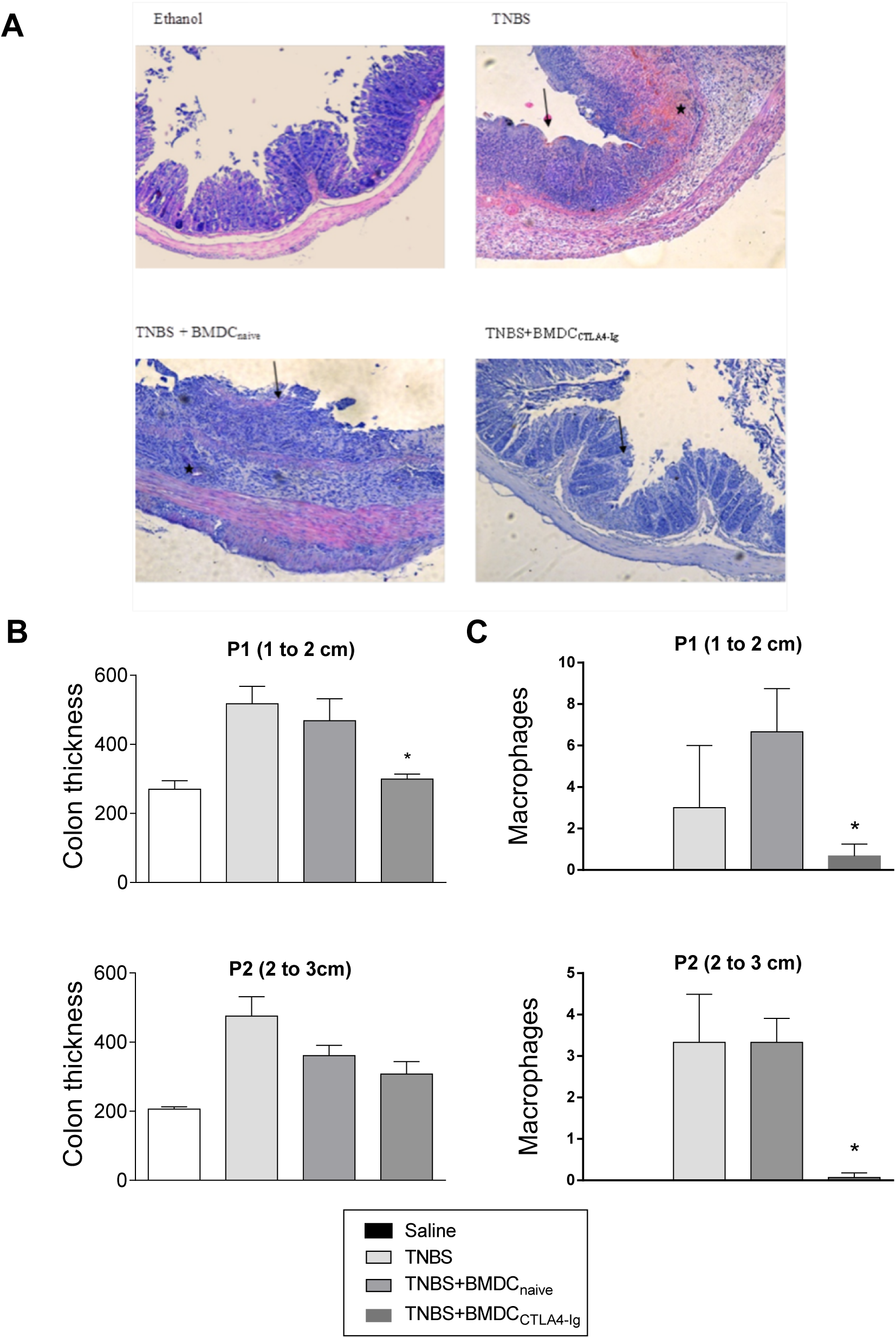

### The effects of adoptive transfer of BMDCs on immune response of colitic mice

The effects of the adoptive transfer of BMDC_CTLA4-Ig_ on the immunological functions of lymphocytes from colitic mice are shown in Figures 4 and 5. The proliferative response of T lymphocytes was significantly lower in cultures of spleen cells from mice pretreated with BMDC_CTLA4-Ig_ than in controls groups (Figure 4A). The frequencies of Treg cells (CD25^+^ Foxp3^+^ T cells) in cultures of spleen cells of mice previously treated with BMDC_CTLA4-Ig_ as well as those receiving TNBS alone were significantly higher than in the other groups, as shown in Figure 4B. The frequency of CD4^+^ T cells producing IFN-γ and IL-17 was significantly lower in the cultures of spleen cells from mice pretreated with BMDC_CTLA4-Ig_, compared to the other groups (Fig 4, C and E). On the other hand, the frequency of CD4^+^ T cells producing IL-10 was higher in the cultures of spleen cells from mice pretreated with BMDC_CTLA4-Ig_ and, interestingly, lower in the cultures of spleen cells from mice pretreated with BMDC_naïve_ compared with the other groups (Figure 4G). No significant differences were observed between the experimental groups in relation to the intracellular labeling of RORc and T-bet transcription factors (Fig 4D, 4F). However, GATA3 factor labeling was higher in spleen cells from mice treated with BMDC_CTLA4-Ig_ (Fig. 4H).

As shown in Figure 5C, IL-4 was present at levels detectable only in spleen cell culture supernatants from mice pretreated with BMDC_CTLA4-Ig_. Significantly elevated IL-10 levels were found in supernatants from splenic cell cultures of mice pretreated with BMDC_naïve_ or BMDC_CTLA4-Ig_, compared to controls (Figure 5E). However, IL-6 levels were also higher in spleen cultures from BMDC_naïve_-pretreated mice compared to the other groups (Figure 5B). Higher IL-9 levels were detected in splenic cell culture supernatants from control mice that received only intrarectal ethanol. In mice treated with BMDC_CTLA4-Ig_ prior to TNBS instillation, however, levels of IL-9 were significantly lower than in all other groups (Figure 5G). There were no significant differences in the levels of IL-2, IL-17, TNF-α and IFN-γ (Figure 5 A, D, F, H, respectively).

**Figure 4:**
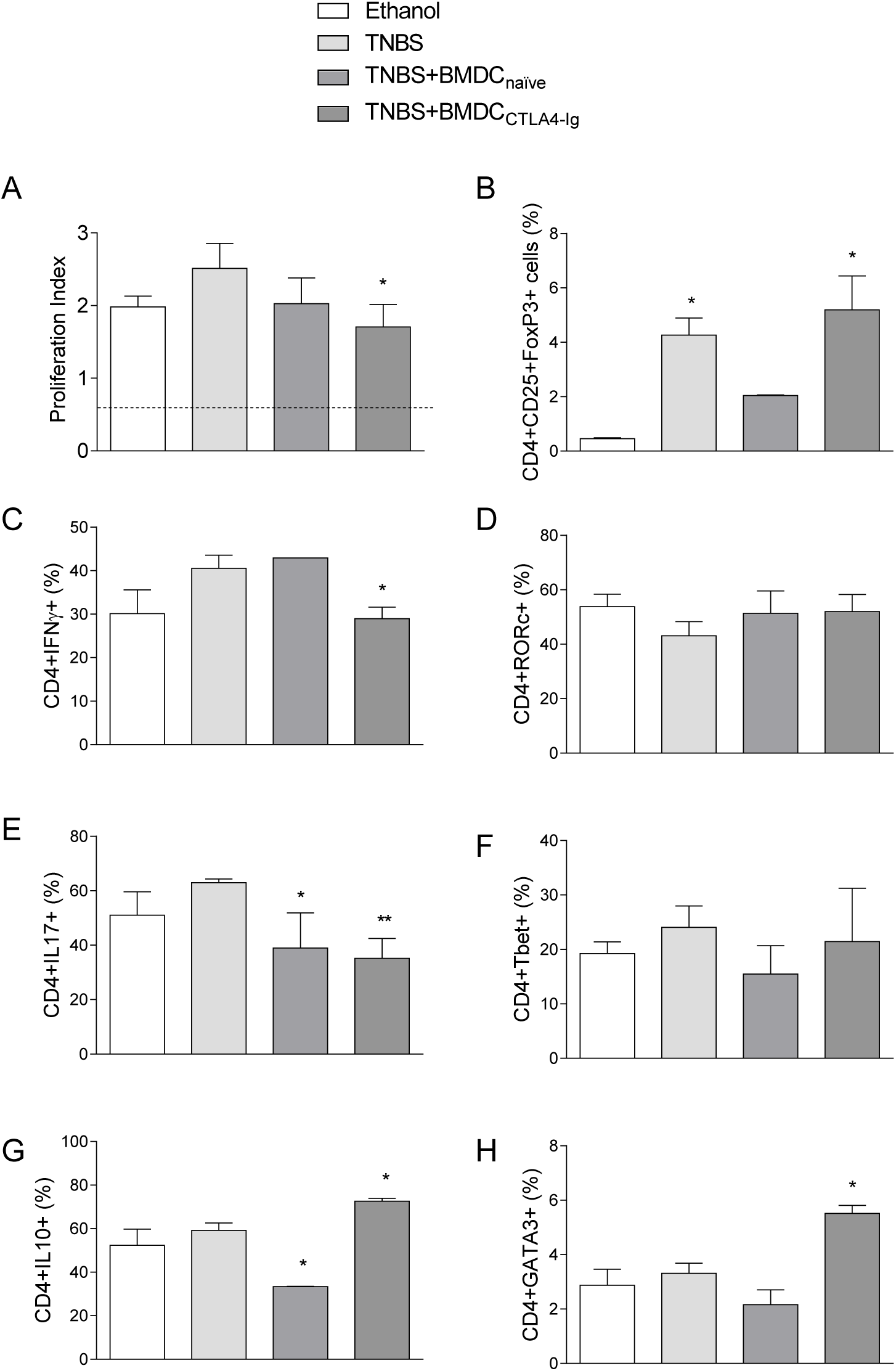

**Figure 5.**
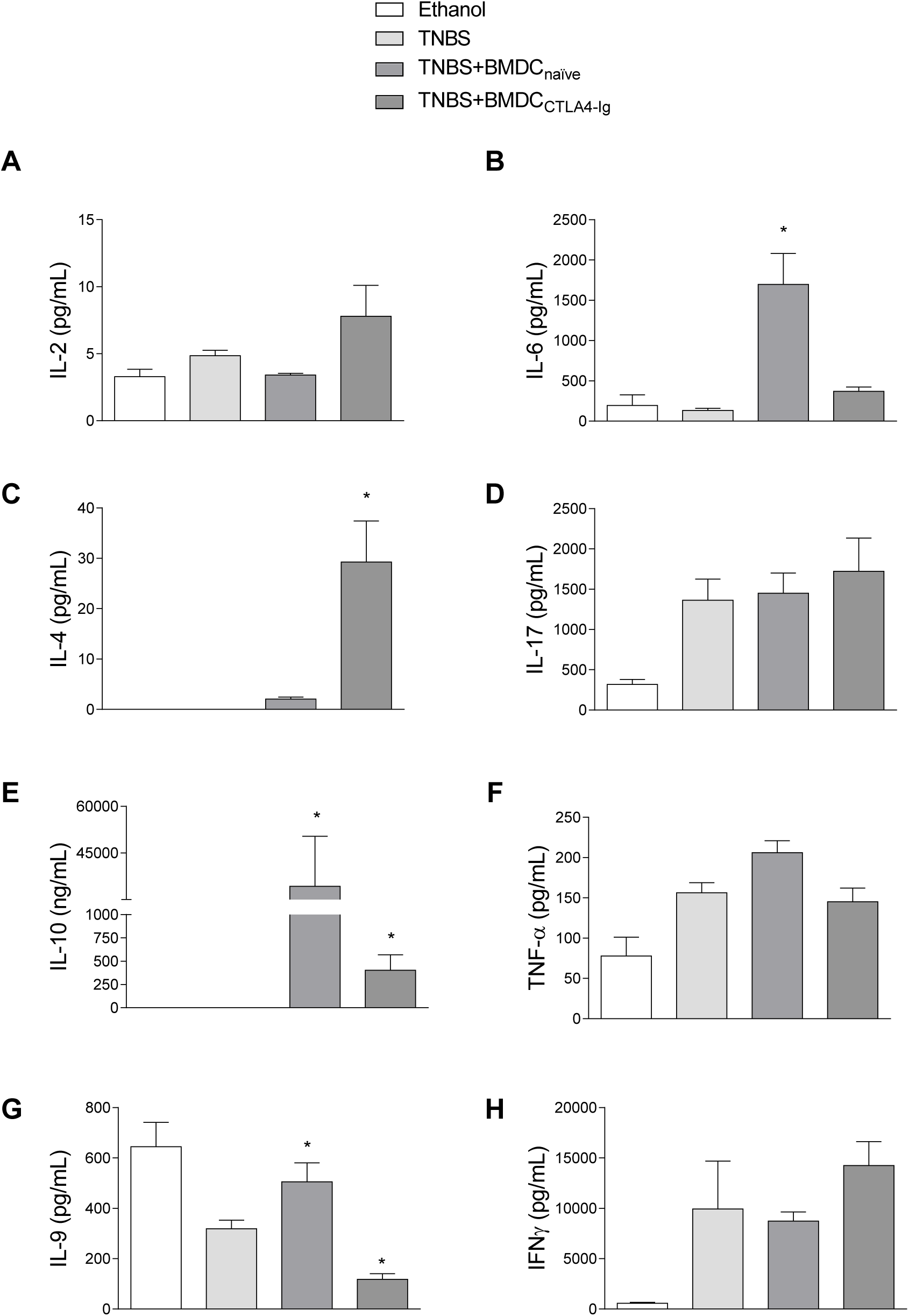

## 4. Discussion

Experimental colitis induced by TNBS instillation is characterized by chronic inflammation of the gastrointestinal tract of mice with features overlapping those seen in inflammatory bowel diseases in humans. Literature data have shown the importance of oral tolerance and treatments with tolerogenic dendritic cell for the reduction of colitis damages In this sense, previous studies of our group have shown that oral tolerance to OVA albumin as well as the adoptive transfer of dendritic cells from OVA-tolerant mice is able to reduce the damage caused by TNBS-induced colitis in syngeneic animals (23,24). We have also shown that flaxseed protein hydrolysates and phenolic fractions were able to ameliorate TNBS-induced colitis in BALB/c mice. Treatments with flaxseed protein fractions reduced inflammation of the intestinal mucosa in TNBS-induced colitis in BALB/c mice, as well as the proliferation of their splenic cells in response to Con-A, the frequency of Th1 and Th17 cells, and the levels of inflammatory cytokines in culture supernatants. In addition, the administration of phenolic compounds from flaxseeds prevented intestinal inflammation and increased the frequency of Treg lymphocytes in splenic cell cultures of BALB/c mice with colitis (34). The present work expanded these findings, demonstrating that the adoptive transfer of bone marrow-derived dendritic cells modulated with CTLA4-Ig, a recombinant mouse protein which binds with both B7-1 and B7-2 molecules, can improve clinical signs of the TNBS-induced colitis in BALB/c mice. It also shows that after transfer of CTLA4-Ig-modulated BMDCs, spleen T lymphocytes from mice with colitis show a more reduced proliferative response to Con-A accompanied by a reduction in the frequency of inflammatory cells secreting IL-17 and IFN-gamma as well as expansion of cells that produce IL-10 in the cultures. Our data showing an improvement in colitis with CTLA4-Ig-modulated BMDCs corroborates with previous data showing that adoptive transfer of DCs modulated with dexamethasone and Vitamin D3 (42,43) or IL-10-modulated DC (44) protects severe combined immune deficient (SCID) mice from weight loss and pathologies associated with wasting diseases and colitis.

CTLA4-Ig is able to selectively modulate T cell activation by binding to CD80/CD86 costimulatory molecules in DCs (9,45). It is already known that direct administration of CTLA4-Ig affects the functioning of DCs through the IDO pathway, promoting a regulatory phenotype and consequently inducing the increase in the population of CD4^+^CD25^+^Foxp3^+^ T cells (9,46,47). In the murine model of arthritis, treatment with CTLA4-Ig was able to reduce the expression of CD80/CD86 molecules on DC and suppressed the inflammatory response associated with the disease (47). Our results did not show significant changes in CD80 and CD86 expression after the BMDCs were treated with CTLA4-Ig. This may be related to the origin of dendritic cells, i.e. differentiated dendritic cells from bone marrow precursors, and to the doses of CTLA4-Ig used in this work. Moreover, we used the recombinant protein to modulate the BMDCs to be transferred adoptively to mice rather than administering it directly to the animals.

It is well known that the instillation of TNBS causes severe changes in the distal portion of the large intestine, due to the inflammatory process triggered by the immune response to the drug. Administration of TNBS to rats, for example, results in increased expression of fibrosis-associated proteins such as phospho-p38, phospho-SMAD2/3, and PPARγ (48). In agreement with previous studies, we observed that the instillation of TNBS caused significant histological changes in the large intestine segments of BALB/c mice, particularly in the P2 segment (2 to 3 cm of the anal sphincter). These changes were characterized by a thickening of the colon and intense inflammatory infiltrate consisting mainly of macrophages.

The literature shows that in the TNBS-induced colitis the adaptive immune response is predominantly Th1 type, characterized by an increase in IFNγ-producing T cells. In protocols for weekly administration of TNBS for six consecutive weeks, an influx of T cells was observed around the third day to two weeks after instillation of the drug, infiltrating the lamina itself and the submucosal layer of the large intestine and supporting chronic colitis (49). In TNBS single-dose protocols such as that used in this study, lymphocyte migration to the lamina propria begins about one week after instillation of the drug (50). Since the animals were euthanized on the fifth day after TNBS administration, this cell type was virtually absent in our histological preparations. Likewise, a reduced number of neutrophils were observed in P1 and P2 preparations since the maximum migration of these cells occurs within the first 48 hours after the instillation of TNBS. As expected, in the time elapsed between administration of TNBS and the euthanasia of the animals for histological analysis, macrophages were the most abundant cells in the inflammatory infiltrate, particularly in the P2 segment of the colon of animals receiving TNBS alone, in a typical hypersensitivity reaction, as described in figures 3.

Our results show that the adoptive transfer of CTLA4-Ig-modulated BMDC was able to significantly prevent colon thickening in the P2 portion of the large intestine as well as the infiltration of macrophages in response to instillation of TNBS. Data from the literature indicate that the intense leukocyte infiltrate in the intestinal mucosa may be responsible for tissue necrosis and changes associated with colitis symptoms (51). Lesions in the colon mucosa may be associated with the release of significant amounts of free radicals, due to the abundance of activated macrophages attracted to the lesion site (52,53). Thus, our results indicate that DCs modulated in vitro with the recombinant CTLA4-Ig protein constitute at least one more natural therapeutic alternative for the treatment of these disorders.

In order to evaluate the influence of the adoptive transfer of CTLA4-Ig-modulated BMDCs on the immune response of TNBS-treated mice, we examined the proliferative responses, the effector CD4^+^ T cell profiles and the release of cytokines in cultures of spleen cells collected on the fifth day after induction of colitis and stimulated in vitro with Con-A. Data presented here (Fig 4) show that spleen cells from animals of all groups proliferated in response to Con-A, but such ability was significantly lower in splenic cells from mice pretreated with CTLA4-Ig-modulated BMDCs.

Examination of effector CD4^+^ T cell populations in splenic cell cultured in the presence of Con-A showed that treatment with BMDC_CTLA4-Ig_ resulted in a significant reduction in the frequency of IL-17^+^ and IFN-γ^+^ cells and in the elevation of CD4^+^ IL-10^+^ and CD4^+^ Foxp3^+^ T cells. Frequency of GATA-3 expressing cells was higher in the splenic cell cultures of mice treated with CTLA4-Ig modulated BMDCs. However, TCD4^+^ cells expressing the Th1/Treg cell associated RORc and T-bet transcription factors did not show significant variations between the different treatments.

Treg cells play a key role in the control of immune responses to autoantigens as well as on those that act upon pathogens, commensals, tumors, and grafts. Such control is exerted by the ability of Treg cells to accumulate in inflamed areas and to adapt to the environment, being particularly critical in tissues repeatedly exposed to the presence of microbes and environmental aggressions such as the gastrointestinal tract and skin (54,55). It has been shown that the canonical Th2 transcription factor GATA3 is selectively expressed in Treg residing in barrier sites including the gastrointestinal tract and the skin, being fundamental to maintain high levels of Foxp3 expression in various polarized or inflammatory settings (56). Corroborating these data, we observed a significantly higher frequency of Treg cells in spleen cell cultures from mice receiving only TNBS and those from mice pretreated with BMDC_CTLA4-Ig_. However, a significant increase in cells expressing both Foxp3 and GATA-3 was observed only in the group that received BMDC_CTLA4-Ig_, indicating its influence in promoting a more efficient control of the inflammatory response induced by TNBS.

Although the frequencies of IL-17 and IFN-γ-secreting T cells were reduced in splenic cell cultures of mice pretreated with BMDC_CTLA4-Ig_, no significant differences were observed in the levels of these cytokines in spleen cell culture supernatants from mice pretreated with either BMDC_CTLA4-Ig_ or BMDC_naïve_. On the other hand, IL-4, whose production is controlled by GATA-3 expression, was detected only in spleen cell cultures of BMDC_CTLA4-Ig_ treated mice. In the spleen cell cultures from mice pretreated with BMDC_naïve_ it was possible to observe the higher levels of IL-6, but it was also the one that presented the highest levels of IL-10, whose production is controlled by the transcription factor Foxp3. Thus, while splenic cell cultures of BMDC_CTLA4-Ig_ pretreated mice had higher levels of IL-4, those of mice pretreated with BMDCnaïve had higher levels of IL-10. The significance of these findings still needs further investigation.

The presence of cells expressing the transcription factor PU.1, a regulator of the development of Th9 cells, has been observed in the intestinal lamina propria of patients with ulcerative colitis and Crohn’s disease (57). Although we did not examine the frequency of Th9 cells in the splenic cell cultures of the different groups studied here, we found that the production of IL-9 in the supernatants of the spleen cell cultures from mice treated with BMDC_CTLA4-Ig_ was significantly more reduced than in other cell cultures, including those from mice that received only the vehicle.

IL-9 is a cytokine that may act differently on Th17 cells or Treg cells, as an inducer or regulator of tissue inflammation. IL-9 associated with TGF-β may drive the differentiation of Th17 cells. In turn, Th17 cells can secrete IL-9, which affects inflammatory response *in vivo*. IL-9 also acts *in vitro* on FoxP3^+^CD4^+^ Treg cells, increasing their suppressive function. This activation occurs by signaling pathways associated with transcription factors STAT3 and STAT5 (58).

Reports show that in addition to cytokines released by Th1 and Th17 cells, IL-9 is also involved in T cell-mediated experimental colitis, promoting mucosal ulceration and chronic inflammation. In this way, Th9 cells represent a potential target for the treatment of chronic intestinal inflammation (22,59).

It is known that acute and chronic intake of alcohol produces sensitive changes in the intestinal mucosa, contributing to the installation or worsening of IBD already installed, both in humans and in experimental models (60). The literature, however, does not report on possible changes due to the use of 50% alcohol as a control of the instillation of TNBS dissolved in this vehicle. However, we have observed that the instillation of 50% ethanol is not as innocuous as its use has resulted in some changes of an inflammatory nature such as elevated TNF-α, IL-9 and IL-17 levels in the corresponding splenic cell cultures, although important differences were observed in the instillation of TNBS/50% ethanol compared to ethanol instillation alone.

The blockade of the CTLA4 molecule is already described as a potential therapy for tumor treatment (61–63). However, studies related to the blockade of this molecule by the direct administration of CTLA4-Ig for the treatment of inflammatory bowel diseases did not present promising results (64). In this context, the use of CTLA4-Ig-modulated dendritic cells, instead of the direct application of this inhibitor, may be a clinical alternative to treat patients with IBD. Studies have shown that the CTLA4-Ig fusion protein affects the functioning of DCs through the IDO pathway, promoting a regulatory phenotype in this population (9,26,47). Dendritic cell therapies for immunomodulation have been presented as a therapeutic option under study, due to the great advance in the use of these cellular populations in the treatments of autoimmune diseases (65,66). Wang and colleagues observed that BMDCs generated from mouse bone marrow and stimulated with GM-CSF have a mature DC cell profile and can be used in antitumor immunity studies. Adherent cells from these cultures have macrophage properties and may be used to induce tolerance, whereas mixed cells may potentiate tolerogenicity or pro-tumorigenic responses. Immature DCs have a strong migration and capture capabilities, while mature DCs activate naïve T cells and express high levels of costimulatory adhesion molecules and cytokines (67).

Taken together, our results allow us to conclude that adoptive transfer of CTLA4-Ig-modulated BMDC improves clinical signs of TNBS-induced colitis. Histological analysis of intestinal segments showed that the adoptive transfer of CTLA4-Ig-modulated BMDC reduced the infiltration of inflammatory cells, particularly macrophages, and improved tissue damage in the colon. Adoptive transfer of CTLA4-Ig-modulated BMDC was also able to alter the immunological profile of activated splenic cells in vitro. Spleen cell culture of CTLA4-Ig-modulated BMDC-pretreated mice showed a reduction in the frequency of CD4^+^ T cells producing IFN-γ and IL-17 and IL-9 secretion, as well as increased frequency of Treg cells and IL-10 production. To our knowledge, this is the first description of the beneficial effects of treatment with CTLA4-Ig modulated BMDC in experimental colitis at the histological and immunological level.

## Data Availability

The research data used to support the findings of this study are included within the article.

---

**Supplementary figure 1.**
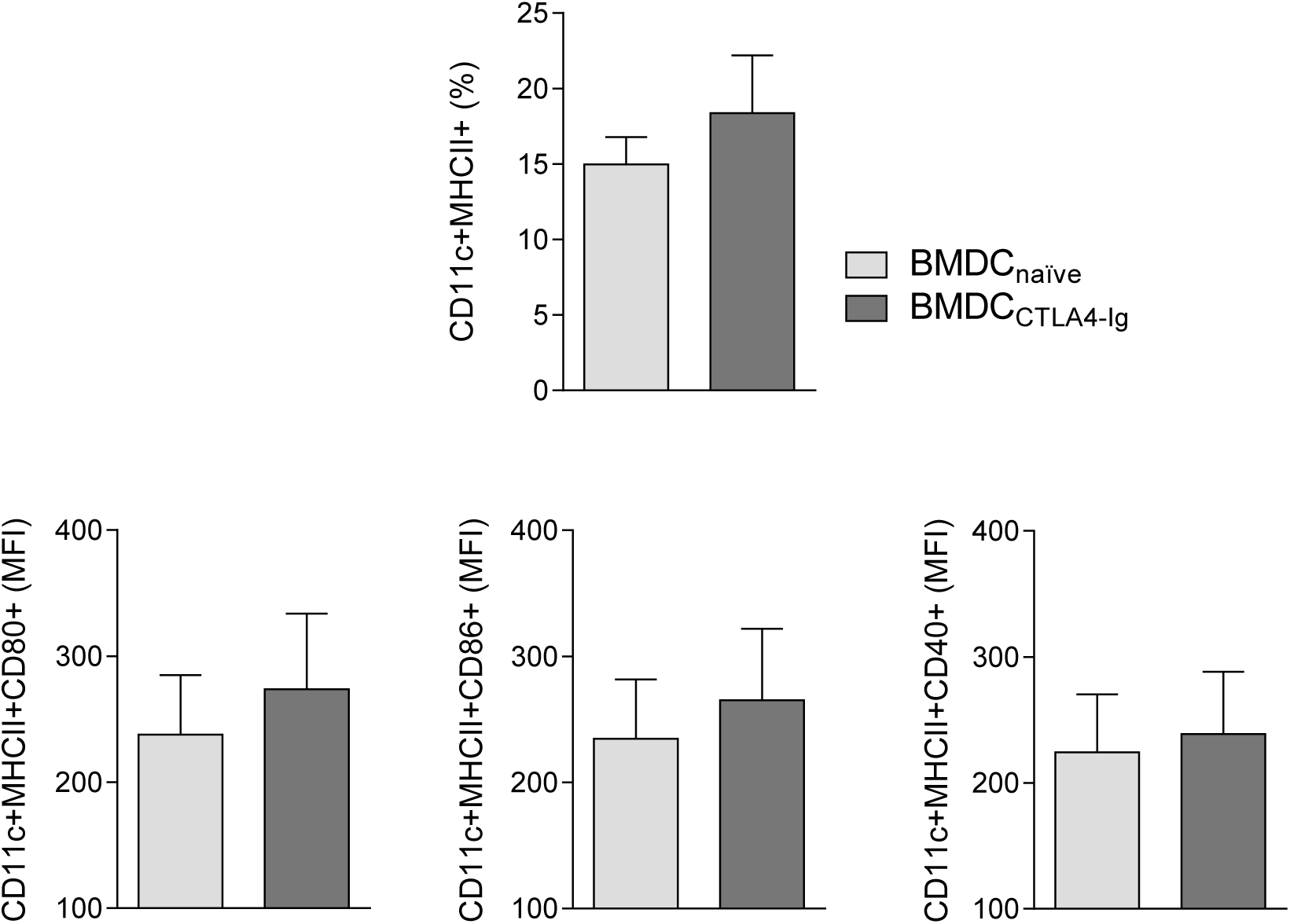

## REFERENCES

1. Metzler B, Burkhart C, Wraith DC. Phenotypic analysis of CTLA-4 and CD28 expression during transient peptide-induced T cell activation in vivo. Int Immunol [Internet]. 1999;11(5):667–75. Available from: https://www.ncbi.nlm.nih.gov/pubmed/10330272

2. Butte MJ, Keir ME, Phamduy TB, Sharpe AH, Freeman GJ. Programmed Death-1 Ligand 1 Interacts Specifically with the B7-1 Costimulatory Molecule to Inhibit T Cell Responses. Immunity [Internet]. 2007;27(1):111–22. Available from: https://www.ncbi.nlm.nih.gov/pmc/articles/PMC2707944/

3. Munn DH, Sharma MD, Mellor AL. Ligation of B7-1/B7-2 by Human CD4+ T Cells Triggers Indoleamine 2,3-Dioxygenase Activity in Dendritic Cells. J Immunol [Internet]. 2004 Mar 19 [cited 2015 Mar 16];172(7):4100–10. Available from: http://www.ncbi.nlm.nih.gov/pubmed/15034022

4. Koorella C, Nair JR, Murray ME, Carlson LM, Watkins SK, Lee KP. Novel regulation of CD80/CD86-induced phosphatidylinositol 3-kinase signaling by NOTCH1 protein in interleukin-6 and indoleamine 2,3-dioxygenase production by dendritic cells. J Biol Chem [Internet]. 2014 Mar 14 [cited 2015 Feb 27];289(11):7747–62. Available from: http://www.pubmedcentral.nih.gov/articlerender.fcgi?artid=3953285&tool=pmcentrez&rendertype=abstract

5. Cutolo M, Soldano S, Montagna P, Sulli A, Seriolo B, Villaggio B, et al. CTLA4-Ig interacts with cultured synovial macrophages from rheumatoid arthritis patients and downregulates cytokine production. Arthritis Res Ther [Internet]. 2009;11(6):R176. Available from: https://www.ncbi.nlm.nih.gov/pmc/articles/PMC3003520/

6. Perez N, Karumuthil-Melethil S, Li R, Prabhakar BS, Holterman MJ, Vasu C. Preferential costimulation by CD80 results in IL-10-dependent TGF-beta1(+)-adaptive regulatory T cell generation. J Immunol [Internet]. 2008 May 15 [cited 2014 Dec 29];180(10):6566–76. Available from: http://www.pubmedcentral.nih.gov/articlerender.fcgi?artid=2435403&tool=pmcentrez&rendertype=abstract

7. Ueda H, Howson JMM, Esposito L, Heward J, Snook H, Chamberlain G, et al. Association of the T-cell regulatory gene CTLA4 with susceptibility to autoimmune disease. Nature [Internet]. 2003 May 29 [cited 2015 Oct 16];423(6939):506–11. Available from: http://www.ncbi.nlm.nih.gov/pubmed/12724780

8. Abram DM, Fernandes LGRL, Ramos Filho AACS, Simioni PUPU, Moitinho Abram D, Fernandes LGRL, et al. The modulation of enzyme indoleamine 2,3-dioxygenase from dendritic cells for the treatment of type 1 diabetes mellitus. Drug Des Devel Ther [Internet]. 2017 Jul [cited 2018 Sep 21];2171–8. Available from: https://www.dovepress.com/the-modulation-of-enzyme-indoleamine-23-dioxygenase-from-dendritic-cel-peer-reviewed-article-DDDT

9. Cutolo M, Nadler SG. Advances in CTLA-4-Ig-mediated modulation of inflammatory cell and immune response activation in rheumatoid arthritis. Autoimmun Rev [Internet]. 2013 May [cited 2015 Feb 18];12(7):758–67. Available from: http://www.ncbi.nlm.nih.gov/pubmed/23340277

10. De Mattos BRR, Garcia MPG, Nogueira JB, Paiatto LNLN, Albuquerque CG, Souza CL, et al. Inflammatory Bowel Disease: An Overview of Immune Mechanisms and Biological Treatments. Mediators Inflamm [Internet]. 2015 Jan 4 [cited 2015 Sep 8];2015:1–11. Available from: http://www.hindawi.com/journals/mi/aa/493012/

11. Corridoni D, Arseneau KO, Cominelli F. Inflammatory bowel disease. Immunol Lett [Internet]. 2014 Oct [cited 2016 Jan 12];161(2):231–5. Available from: http://www.pubmedcentral.nih.gov/articlerender.fcgi?artid=4401421&tool=pmcentrez&rendertype=abstract

12. Aujnarain A, Mack DR, Benchimol EI. The role of the environment in the development of pediatric inflammatory bowel disease. Curr Gastroenterol Rep [Internet]. 2013 Jun [cited 2014 Dec 12];15(6):326. Available from: http://www.ncbi.nlm.nih.gov/pubmed/23640032

13. Tontini GE. Differential diagnosis in inflammatory bowel disease colitis: State of the art and future perspectives. World J Gastroenterol [Internet]. 2015 Jan 7 [cited 2015 Jan 10];21(1):21. Available from: http://www.pubmedcentral.nih.gov/articlerender.fcgi?artid=4284336&tool=pmcentrez&rendertype=abstract

14. Khan A, Fu H, Tan LA, Harper JE, Beutelspacher SC, Larkin DFP, et al. Dendritic cell modification as a route to inhibiting corneal graft rejection by the indirect pathway of allorecognition. Eur J Immunol [Internet]. 2013 Mar [cited 2015 Jun 12];43(3):734–46. Available from: http://www.pubmedcentral.nih.gov/articlerender.fcgi?artid=3615172&tool=pmcentrez&rendertype=abstract

15. Najafian N, Sayegh MH. CTLA4-Ig: a novel immunosuppressive agent. Expert Opin Investig Drugs [Internet]. 2000;9(9):2147–57. Available from: http://www.ncbi.nlm.nih.gov/pubmed/11060799

16. Steurer W, Nickerson PW, Steele AW, Steiger J, Zheng XX, Strom TB. Ex vivo coating of islet cell allografts with murine CTLA4/Fc promotes graft tolerance. J Immunol [Internet]. 1995;155(3):1165–74. Available from: http://www.ncbi.nlm.nih.gov/pubmed/7543517

17. Yang DF, Qiu WH, Zhu HF, Lei P, Wen X, Dai H, et al. CTLA4-Ig-modified dendritic cells inhibit lymphocyte-mediated alloimmune responses and prolong the islet graft survival in mice. Transpl Immunol [Internet]. 2008;19(3– 4):197–201. Available from: http://www.ncbi.nlm.nih.gov/pubmed/18667318%5Cnhttp://ac.els-cdn.com/S0966327408000294/1-s2.0-S0966327408000294-main.pdf?_tid=1f04f6c0-df97-11e3-befe-00000aab0f6b&acdnat=1400532771_9f1bf87c9da1bf8eeaae21083b16bd8f

18. Gilson CR, Milas Z, Gangappa S, Hollenbaugh D, Pearson TC, Ford ML, et al. Anti-CD40 monoclonal antibody synergizes with CTLA4-Ig in promoting long-term graft survival in murine models of transplantation. J Immunol [Internet]. 2009 Aug 1 [cited 2015 Jun 16];183(3):1625–35. Available from: http://www.pubmedcentral.nih.gov/articlerender.fcgi?artid=2828346&tool=pmcentrez&rendertype=abstract

19. … ASITS in D, 2019 undefined. Disease Interception in Autoimmune Diseases: From a Conceptual Framework to Practical Implementation. books.google.com [Internet]. [cited 2019 Feb 20]; Available from: https://books.google.com.br/books?hl=pt-BR&lr=&id=uNyGDwAAQBAJ&oi=fnd&pg=PA1&dq=ctla4-ig+tolerance+colitis&ots=NeTRlZ1bvJ&sig=3wf0H2u9tjKcXGtdgXrPzM9xPyI

20. Elson CO, Sartor RB, Tennyson GS, Riddell Ii RH, Gonçalves CCM, Hernandes L, et al. Insights from advances in research of chemically induced experimental models of human inflammatory bowel disease. World J Gastroenterol. 1995;13(42):5581–93.

21. Oh SY, Cho K-A, Kang JL, Kim KH, Woo S-Y. Comparison of experimental mouse models of inflammatory bowel disease. Int J Mol Med [Internet]. 2014 Feb [cited 2016 Jan 6];33(2):333–40. Available from: http://www.ncbi.nlm.nih.gov/pubmed/24285285

22. Gerlach K, Hwang Y, Nikolaev A, Atreya R, Dornhoff H, Steiner S, et al. TH9 cells that express the transcription factor PU.1 drive T cell-mediated colitis via IL-9 receptor signaling in intestinal epithelial cells. Nat Immunol [Internet]. 2014 Jul 8 [cited 2014 Dec 3];15(7):676–86. Available from: http://www.ncbi.nlm.nih.gov/pubmed/24908389

23. Paiatto LN, Silva FGD, Simioni PU, Yamada ÁT, Tamashiro WMSC, Simioni PU. Adoptive transfer of dendritic cells expressing CD11c reduces the immunological response associated with experimental colitis in BALB / c mice. Chamaillard M, editor. PLoS One [Internet]. 2018 May 8 [cited 2018 Sep 21];13(5):1–15. Available from: http://dx.plos.org/10.1371/journal.pone.0196994

24. Paiatto LNLN, Silva FGDFGDFGD, Bier J, Brochetto-Braga MR, Yamada ÁTÁT, Tamashiro WMSCWMSCSC, et al. Oral tolerance induced by OVA intake ameliorates TNBS-induced colitis in Mice. Ashour HM, editor. PLoS One [Internet]. 2017 Jan 18 [cited 2017 Jan 24];12(1):e0170205. Available from: http://dx.plos.org/10.1371/journal.pone.0170205

25. Matteoli G, Mazzini E, Iliev ID, Mileti E, Fallarino F, Puccetti P, et al. Gut CD103+ dendritic cells express indoleamine 2,3-dioxygenase which influences T regulatory/T effector cell balance and oral tolerance induction. Gut [Internet]. 2010 May [cited 2013 Jun 7];59(5):595–604. Available from: http://gut.bmj.com/cgi/doi/10.1136/gut.2009.185108

26. Mayer L, Kaser A, Blumberg RS. Dead on arrival: understanding the failure of CTLA4-immunoglobulin therapy in inflammatory bowel disease. Gastroenterology [Internet]. 2012 Jul [cited 2014 Dec 12];143(1):13–7. Available from: http://www.pubmedcentral.nih.gov/articlerender.fcgi?artid=3392152&tool=pmcentrez&rendertype=abstract

27. Qualls JE, Tuna H, Kaplan AM, Cohen D a. Suppression of experimental colitis in mice by CD11c+ dendritic cells. Inflamm Bowel Dis [Internet]. 2009 Feb [cited 2014 Nov 24];15(2):236–47. Available from: http://www.ncbi.nlm.nih.gov/pubmed/18839426

28. Linsley PS, Brady W, Urnes M, Grosmaire LS, Damle NK, Ledbetter JA. CTLA-4 is a second receptor for the B cell activation antigen B7. J Exp Med [Internet]. 1991;174(3):561–9. Available from: http://www.ncbi.nlm.nih.gov/pubmed/1714933%0Ahttp://www.pubmedcentral.nih.gov/articlerender.fcgi?artid=PMC2118936

29. Simioni PUPUPU, Fernandes LGRLGRL, Gabriel DLL, Tamashiro WMSCMSC. Effect of aging and oral tolerance on dendritic cell function. Brazilian J … [Internet]. 2010 Jan [cited 2015 Mar 27];43(1):68–76. Available from: http://www.ncbi.nlm.nih.gov/pubmed/19967261

30. Simioni PUPUU, Fernandes LGRGRLG, Tamashiro WMWMSCMWM. Downregulation of L-arginine metabolism in dendritic cells induces tolerance to exogenous antigen. Int J Immunopathol Pharmacol [Internet]. 2017 Nov 30 [cited 2016 Dec 2];30(1):44–57. Available from: https://www.ncbi.nlm.nih.gov/pubmed/27903843

31. Lutz MB, Kukutsch N a., Menges M, Rössner S, Schuler G, Rößner S, et al. Culture of bone marrow cells in GM-CSF plus high doses of lipopolysaccharide generates exclusively immature dendritic cells which induce alloantigen-specific CD4 T cell anergy in vitro. Eur J Immunol [Internet]. 2000 Apr [cited 2015 Feb 17];30(4):1048–52. Available from: http://www.ncbi.nlm.nih.gov/pubmed/10760792

32. Simioni PUPUU, Fernandes LGRGRLG, Tamashiro WMSCMWM, Lutz MB, Kukutsch N a., Menges M, et al. An advanced culture method for generating large quantities of highly pure dendritic cells from mouse bone marrow. Eur J Immunol [Internet]. 2017 Nov 1 [cited 2015 Feb 6];30(1):1048–52. Available from: http://www.ncbi.nlm.nih.gov/pubmed/10037236

33. Lutz MB, Kukutsch N, Ogilvie AL, Rössner S, Koch F, Romani N, et al. An advanced culture method for generating large quantities of highly pure dendritic cells from mouse bone marrow. J Immunol Methods [Internet]. 1999 Feb 1 [cited 2015 Feb 6];223(1):77–92. Available from: http://www.ncbi.nlm.nih.gov/pubmed/10037236

34. E Silva FGD, Paiatto LN, Yamada AT, Netto FM, Simioni PU, Tamashiro WMSC. Intake of Protein Hydrolysates and Phenolic Fractions Isolated from Flaxseed Ameliorates TNBS-Induced Colitis. Mol Nutr Food Res [Internet]. 2018 Aug 9 [cited 2018 Aug 16];e1800088. Available from: http://doi.wiley.com/10.1002/mnfr.201800088

35. Masterson JC, McNamee EN, Fillon SA, Hosford L, Harris R, Fernando SD, et al. Eosinophil-mediated signalling attenuates inflammatory responses in experimental colitis. Gut [Internet]. 2014 Sep 10 [cited 2015 Mar 9];gutjnl-2014-306998-. Available from: http://gut.bmj.com/content/early/2014/09/10/gutjnl-2014-306998.long

36. Zingarelli B, Szabó C, Salzman AL. Reduced oxidative and nitrosative damage in murine experimental colitis in the absence of inducible nitric oxide synthase. Gut [Internet]. 1999 Aug [cited 2015 Mar 9];45(2):199–209. Available from: http://www.pubmedcentral.nih.gov/articlerender.fcgi?artid=1727621&tool=pmcentrez&rendertype=abstract

37. van der Marel S, Majowicz A, Kwikkers K, van Logtenstein R, te Velde AA, De Groot AS, et al. Adeno-associated virus mediated delivery of Tregitope 167 ameliorates experimental colitis. World J Gastroenterol [Internet]. 2012 Aug 28 [cited 2015 Feb 18];18(32):4288–99. Available from: http://www.pubmedcentral.nih.gov/articlerender.fcgi?artid=3436043&tool=pmcentrez&rendertype=abstract

38. Shin J-S, Cho E-J, Choi H-E, Seo J-H, An H-J, Park H-J, et al. Anti-inflammatory effect of a standardized triterpenoid-rich fraction isolated from Rubus coreanus on dextran sodium sulfate-induced acute colitis in mice and LPS-induced macrophages. J Ethnopharmacol [Internet]. 2014 Dec 2 [cited 2015 Feb 18];158 Pt A:291–300. Available from: http://www.ncbi.nlm.nih.gov/pubmed/25446582

39. Neurath MF, Fuss I, Kelsall BL, Stüber E, Strober W. Antibodies to interleukin 12 abrogate established experimental colitis in mice. J Exp Med [Internet]. 1995 Nov 1 [cited 2016 Dec 2];182(5):1281–90. Available from: http://www.ncbi.nlm.nih.gov/pubmed/7595199

40. Neurath MF, Fuss I, Kelsall BL, Presky DH, Waegell W, Strober W. Experimental granulomatous colitis in mice is abrogated by induction of TGF-beta-mediated oral tolerance. J Exp Med [Internet]. 1996 Jun 1 [cited 2015 Feb 18];183(6):2605–16. Available from: http://www.ncbi.nlm.nih.gov/pubmed/8676081

41. Thomé R, Fernandes LGRLGRL, Mineiro MFMFM, Simioni PUPU, Joazeiro PPPP, Tamashiro WM da SCWMSC. Oral tolerance and OVA-induced tolerogenic dendritic cells reduce the severity of collagen/ovalbumin-induced arthritis in mice. Cell Immunol [Internet]. 2012 Nov [cited 2015 Feb 5];280(1):113–23. Available from: http://linkinghub.elsevier.com/retrieve/pii/S0008874912002201

42. Horton C, Shanmugarajah K, Fairchild PJ. Harnessing the properties of dendritic cells in the pursuit of immunological tolerance [Internet]. Vol. 40, Biomedical Journal. 2017. p. 80–93. Available from: https://www.ncbi.nlm.nih.gov/pmc/articles/PMC6138597/

43. Pedersen AE, Schmidt EGW, Gad M, Poulsen SS, Claesson MH. Dexamethasone/1α-25-dihydroxyvitamin D3-treated dendritic cells suppress colitis in the SCID T-cell transfer model. Immunology [Internet]. 2009 Jul [cited 2016 Dec 2];127(3):354–64. Available from: http://www.ncbi.nlm.nih.gov/pubmed/19019085

44. Pedersen AE, Gad M, Kristensen NN, Haase C, Nielsen CH, Claesson MH. Tolerogenic dendritic cells pulsed with enterobacterial extract suppress development of colitis in the severe combined immunodeficiency transfer model. Immunology [Internet]. 2007 Aug [cited 2013 Jul 24];121(4):526–32. Available from: http://www.ncbi.nlm.nih.gov/pubmed/17428312

45. Rochman Y, Yukawa M, Kartashov A V, Barski A. Functional characterization of human T cell hyporesponsiveness induced by CTLA4-Ig. PLoS One [Internet]. 2015 Jan [cited 2015 Jun 15];10(4):e0122198. Available from: http://www.pubmedcentral.nih.gov/articlerender.fcgi?artid=4393265&tool=pmcentrez&rendertype=abstract

46. Mayer E, Hölzl M, Ahmadi S, Dillinger B, Pilat N, Fuchs D, et al. CTLA4-Ig immunosuppressive activity at the level of dendritic cell/T cell crosstalk. Int Immunopharmacol [Internet]. 2013 Mar [cited 2013 Jun 23];15(3):638–45. Available from: http://dx.doi.org/10.1016/j.intimp.2013.02.007

47. Ko HJ, Cho M La, Lee SY, Oh HJ, Heo YJ, Moon YM, et al. CTLA4-Ig modifies dendritic cells from mice with collagen-induced arthritis to increase the CD4+CD25+Foxp3+ regulatory T cell population. J Autoimmun. 2010;34(2):111–20.

48. Loeuillard E, Bertrand J, Herranen A, Melchior C, Guérin C, Coëffier M, et al. 2,4,6-trinitrobenzene sulfonic acid-induced chronic colitis with fibrosis and modulation of TGF-β1 signaling. World J Gastroenterol [Internet]. 2014 Dec 28 [cited 2016 Jan 5];20(48):18207–15. Available from: http://www.pubmedcentral.nih.gov/articlerender.fcgi?artid=4277958&tool=pmcentrez&rendertype=abstract

49. Wu F, Chakravarti S. Differential Expression of Inflammatory and Fibrogenic Genes and Their Regulation by NF-B Inhibition in a Mouse Model of Chronic Colitis. J Immunol [Internet]. 2007 Nov 2 [cited 2016 Jan 5];179(10):6988–7000. Available from: http://www.jimmunol.org/content/179/10/6988.full

50. Miura S, Hokari R, Tsuzuki Y. Mucosal immunity in gut and lymphoid cell trafficking. Ann Vasc Dis [Internet]. 2012 Jan [cited 2016 Jan 7];5(3):275–81. Available from: http://www.pubmedcentral.nih.gov/articlerender.fcgi?artid=3595844&tool=pmcentrez&rendertype=abstract

51. Guo X, Wang WP, Ko JKS, Cho CH. Involvement of neutrophils and free radicals in the potentiating effects of passive cigarette smoking on inflammatory bowel disease in rats. Gastroenterology [Internet]. 1999 Oct 10 [cited 2016 Jan 5];117(4):884–92. Available from: http://www.gastrojournal.org/article/S0016508599703471/fulltext

52. Seo HG, Takata I, Nakamura M, Tatsumi H, Suzuki K, Fujii J, et al. Induction of nitric oxide synthase and concomitant suppression of superoxide dismutases in experimental colitis in rats. Arch Biochem Biophys [Internet]. 1995 Dec 1 [cited 2016 Jan 5];324(1):41–7. Available from: http://www.ncbi.nlm.nih.gov/pubmed/7503557

53. Yue G, Lai PS, Yin K, Sun FF, Nagele RG, Liu X, et al. Colon epithelial cell death in 2,4,6-trinitrobenzenesulfonic acid-induced colitis is associated with increased inducible nitric-oxide synthase expression and peroxynitrite production. J Pharmacol Exp Ther [Internet]. 2001 Jun [cited 2016 Jan 5];297(3):915–25. Available from: http://www.ncbi.nlm.nih.gov/pubmed/11356911

54. Sakaguchi S. Naturally arising CD4+ regulatory t cells for immunologic self-tolerance and negative control of immune responses. Annu Rev Immunol [Internet]. 2004;22:531–62. Available from: http://arjournals.annualreviews.org/doi/full/10.1146/annurev.immunol.21.120601.141122?url_ver=Z39.88-2003&rfr_id=ori:rid:crossref.org&rfr_dat=cr_pub%3Dpubmed

55. Campbell DJ, Koch MA. Phenotypical and functional specialization of FOXP3+ regulatory T cells. Nat Rev Immunol [Internet]. 2011;11(2):119–30. Available from: http://www.nature.com/doifinder/10.1038/nri2916

56. Wohlfert EEA, Grainger JRJ, Bouladoux N, Konkel JE, Oldenhove G, Ribeiro CH, et al. GATA3 controls Foxp3+ regulatory T cell fate during inflammation in mice. J Clin … [Internet]. 2011;121(11):4503–15. Available from: http://www.ncbi.nlm.nih.gov/pmc/articles/PMC3204837/

57. Kaplan MH. Th9 cells: Differentiation and disease. Immunol Rev [Internet]. 2013;252(1):104–15. Available from: https://onlinelibrary.wiley.com/doi/full/10.1111/imr.12028?sid=nlm%3Apubmed

58. Elyaman W, Bradshaw EM, Uyttenhove C, Dardalhon V, Awasthi A, Imitola J, et al. IL-9 induces differentiation of TH17 cells and enhances function of FoxP3+ natural regulatory T cells. Proc Natl Acad Sci U S A [Internet]. 2009 Aug 4 [cited 2016 Aug 5];106(31):12885–90. Available from: http://www.ncbi.nlm.nih.gov/pubmed/19433802

59. Gerlach K, McKenzie AN, Neurath MF, Weigmann B. IL-9 regulates intestinal barrier function in experimental T cell-mediated colitis. Tissue barriers [Internet]. 2015 Apr 3 [cited 2016 Sep 29];3(1–2):e983777. Available from: http://www.ncbi.nlm.nih.gov/pubmed/25838986

60. Bishehsari F, Magno E, Swanson G, Desai V, Voigt RM, Forsyth CB, et al. Alcohol and Gut-Derived Inflammation. Alcohol Res Curr Rev [Internet]. 2017;38(2):e1–9. Available from: https://www.arcr.niaaa.nih.gov/arcr382/article01.pdf

61. Berman D, Parker SM, Siegel J, Chasalow SD, Weber J, Galbraith S, et al. Blockade of cytotoxic T-lymphocyte antigen-4 by ipilimumab results in dysregulation of gastrointestinal immunity in patients with advanced melanoma. Cancer Immun [Internet]. 2010 Jan [cited 2016 Jan 5];10:11. Available from: http://www.pubmedcentral.nih.gov/articlerender.fcgi?artid=2999944&tool=pmcentrez&rendertype=abstract

62. Daud A. Current and Emerging Perspectives on Immunotherapy for Melanoma. Semin Oncol [Internet]. 2015 Dec [cited 2015 Nov 27];42:S3–11. Available from: http://www.ncbi.nlm.nih.gov/pubmed/26598057

63. Quirk SK, Shure AK, Agrawal DK. Immune-mediated adverse events of anticytotoxic T lymphocyte-associated antigen 4 antibody therapy in metastatic melanoma. Transl Res [Internet]. 2015 Nov [cited 2016 Jan 5];166(5):412–24. Available from: http://www.ncbi.nlm.nih.gov/pubmed/26118951

64. Kawalec P, Mikrut A, Łopuch S. Systematic review of the effectiveness of biological therapy for active moderate to severe ulcerative colitis. J Gastroenterol Hepatol [Internet]. 2014 Jun [cited 2016 Jan 6];29(6):1159–70. Available from: http://doi.wiley.com/10.1111/jgh.12563

65. Cools N, Petrizzo A, Smits E, Buonaguro FM, Tornesello ML, Berneman Z, et al. Dendritic cells in the pathogenesis and treatment of human diseases: a Janus Bifrons? Immunotherapy [Internet]. 2011 Oct [cited 2015 Feb 17];3(10):1203–22. Available from: http://www.ncbi.nlm.nih.gov/pubmed/21995572

66. Harden JL, Egilmez NK. Indoleamine 2,3-Dioxygenase and Dendritic Cell Tolerogenicity [Internet]. Immunological Investigations NIH Public Access; Aug 27, 2012 p. 738–64. Available from: http://www.ncbi.nlm.nih.gov/pubmed/23017144

67. Wang J, Dai X, Hsu C, Ming C, He Y, Zhang J, et al. Discrimination of the heterogeneity of bone marrow-derived dendritic cells. Mol Med Rep. 2017;16(5):6787–93.

